# Raising the stakes: Loss of efflux-pump regulation decreases meropenem susceptibility in *Burkholderia pseudomallei*

**DOI:** 10.1101/205070

**Authors:** Derek S. Sarovich, Jessica R. Webb, Matthew C. Pitman, Linda T. Viberg, Mark Mayo, Robert W. Baird, Jennifer M. Robson, Bart J. Currie, Erin P. Price

## Abstract

*Burkholderia pseudomallei*, the causative agent of the high-mortality disease melioidosis, is a Gram-negative bacterium that is naturally resistant to many antibiotics. There is no vaccine for melioidosis, and effective eradication is reliant on biphasic and prolonged antibiotic administration. The carbapenem drug, meropenem, is the current gold-standard option for treating severe melioidosis. Intrinsic *B. pseudomallei* resistance towards meropenem has not yet been documented; however, resistance could conceivably develop over the course of infection, leading to prolonged sepsis and treatment failure. Here, we document 11 melioidosis cases in which *B. pseudomallei* isolates developed decreased susceptibility towards meropenem during treatment, including two cases not treated with this antibiotic. Meropenem minimum inhibitory concentrations increased over time from 0.5-0.75 to 3-8 μg/mL. Using comparative genomics, we identified multiple mutations affecting multidrug resistance-nodulation-division (RND) efflux pump regulators, leading to over-expression of their corresponding pumps. The most commonly affected pump was AmrAB-OprA, although alterations in the local regulators of BpeEF-OprC or BpeAB-OprB were observed in three cases. This study confirms the role of RND efflux pumps in decreased meropenem susceptibility in *B. pseudomallei*. Further, we document two concerning examples of severe melioidosis where the reduced treatment efficacy of meropenem was associated with a fatal outcome.

**Significance Statement:** The bacterium *Burkholderia pseudomallei*, which causes the often-fatal tropical disease melioidosis, is difficult to eradicate. Due to high levels of intrinsic antibiotic resistance, only a handful of antibiotics are effective against this pathogen. One of these, meropenem, is commonly used in the treatment of melioidosis patients who are unresponsive to other treatments or are critically ill. Here, we describe 11 melioidosis cases whereby patients exhibited prolonged or repeated infections that were associated with the development of decreased meropenem susceptibility. We identified the molecular basis for this decreased susceptibility in latter *B. pseudomallei* isolates obtained from these patients, and functionally confirmed the mechanism conferring this phenotype. Our findings have important ramifications for the diagnosis, treatment and management of life-threatening melioidosis cases.

## Introduction

*Burkholderia pseudomallei* is the causative agent of the potentially deadly disease melioidosis, and is one of the most intrinsically drug-resistant bacterial pathogens (1). As there is currently no effective vaccine for melioidosis, eradiation relies on a lengthy antibiotic regimen (2, 3). Even with appropriate antibiotics, rapid diagnosis and high standards of healthcare, case fatality rates remain at 10% in northern Australia and 40% in Southeast Asia (4, 5), the two regions with the highest reported melioidosis rates (6, 7). *B. pseudomallei* is a significant yet underreported cause of morbidity and mortality in tropical and subtropical regions, with recent modeling suggesting that the global melioidosis mortality rate may be similar to that of measles at ~89,000 deaths per year and substantially higher than that from dengue (7).

Melioidosis treatment is biphasic, comprising an initial intensive intravenous (IV) phase of a minimum of 10-14 days, followed by a prolonged oral eradication phase lasting up to six months. The third-generation cephalosporin antibiotic, ceftazidime, is generally used preferentially in the IV phase. The carbapenems imipenem and meropenem have the lowest minimum inhibitory concentrations (MICs) against *B. pseudomallei*, and *in vitro* time-kill studies have shown that carbapenems perform better against *B. pseudomallei* than ceftazidime (8, 9). Meropenem is recommended for patients who present with life-threatening sepsis requiring intensive care therapy, although there is no clinical evidence that ceftazidime is inferior to meropenem for patients with melioidosis who are not critically ill and ceftazidime remains the drug of choice for initial therapy (2, 3). Importantly, primary resistance towards meropenem has not yet been documented. During the eradication phase of treatment, the preferred treatment is trimethoprim-sulfamethoxazole (TMP/SMX), with amoxicillin-clavulanate or doxycycline used as alternatives when required.

Due to limited treatment options, the emergence of antibiotic resistance during infection, although rare, is a significant event that can lead to treatment failure and subsequent patient death. Unlike many other infectious diseases, melioidosis is largely non-communicable, with almost all infections acquired following contact with *B. pseudomallei*-contaminated soil or water (1). As a result, resistance or decreased susceptibility to clinically-relevant antibiotics emerges as an independent event during treatment rather than by environmental influences or transmission of an antibiotic-resistant strain from another individual or animal (10).

The mechanisms that *B. pseudomallei* evolves to evade antibiotics during an infection are myriad and multifactorial. Unlike many other species, all mechanisms occur as chromosomal alterations; acquisition of extrachromosomal elements containing antibiotic resistance genes has not yet been documented in *B. pseudomallei*. Resistance to ceftazidime can occur by up-regulation or alteration of the β-lactamase PenA (encoded by *penA*; (11–16)) or via the loss of penicillin-binding protein 3 (17). Point mutations in *penA* have recently been shown to lead to imipenem resistance, a carbapenem drug similar to meropenem (18), and missense and promoter mutations affecting *penA* confer resistance towards amoxicillin-clavulanate (12, 15). Up-regulation of multi-drug efflux pumps can also contribute to antibiotic resistance in *B. pseudomallei*. There are three characterized pumps known to confer resistance (AmrAB-OprA, BpeEF-OprC or BpeAB-OprB) and up-regulation of one or more of these pumps in combination with mutations in methylation or metabolism pathways lead to doxycycline or TMP/SMX resistance, respectively (14, 19, 20).

We have recently sequenced to closure, or close-to-closure, isogenic *B. pseudomallei* strain pairs from three melioidosis cases where decreased susceptibility towards meropenem was observed (21). In the current study, we identified the mutations leading to decreased meropenem susceptibility in these pairs using whole-genome sequencing (WGS) and comparative genomics and functionally characterized the mechanism using allelic exchange and quantitative reverse transcription PCR (RT-qPCR). We then reviewed ~1,000 melioidosis cases enrolled in the Darwin Prospective Melioidosis Study (DPMS) (5), an ongoing study that has documented all cases in the endemic “Top End” region of the Northern Territory, Australia, since October 1989. The intent was to identify DPMS patients who experienced difficulties in clearing their infections, pointing towards the possible emergence of *B. pseudomallei* strains with decreased meropenem susceptibility. Patients included those with persistent culture positivity or with repeated recrudescent or relapsing infections, irrespective of whether they were treated with meropenem. Where available, longitudinal isolates were examined to identify the emergence of decreased meropenem susceptibility over time. In all cases where decreased susceptibility was observed, we used comparative genomics to identify the mechanisms underpinning this altered phenotype.

## Results

### Isolates with decreased meropenem susceptibilities have an altered AmrR efflux pump regulator

We have recently identified three DPMS patients (P608, P726 and P797) with severe melioidosis and prolonged blood culture-positivity who harbored *B. pseudomallei* strains exhibiting decreased meropenem susceptibility following meropenem treatment (21). The treatment and admission history for these patients is detailed in Figure S1. P608, P726 and P797 were repeatedly, but not continuously, blood-culture positive for 64, 89, and 454 days, respectively. P726 and P797 ultimately died whereas P608 eventually cleared their infection. The difficulty in eradicating these infections despite prolonged and aggressive antibiotic therapy prompted us to investigate *B. pseudomallei* MICs to the clinically relevant antibiotics amoxicillin-clavulanate, ceftazidime, doxycycline, meropenem and TMP/SMX. All initial isolates were sensitive to meropenem (MICs = 0.5-0.75 μg/mL) whereas the latter isolates had increased MICs that ranged from 3 to 8 μg/mL (Figure 1; Table 1). P608 and P797 isolates had also developed resistance towards TMP/SMX at 24/456 and 4/76 μg/mL, respectively. All isolates remained sensitive to amoxicillin-clavulanate, ceftazidime and doxycycline.

**Figure 1:**
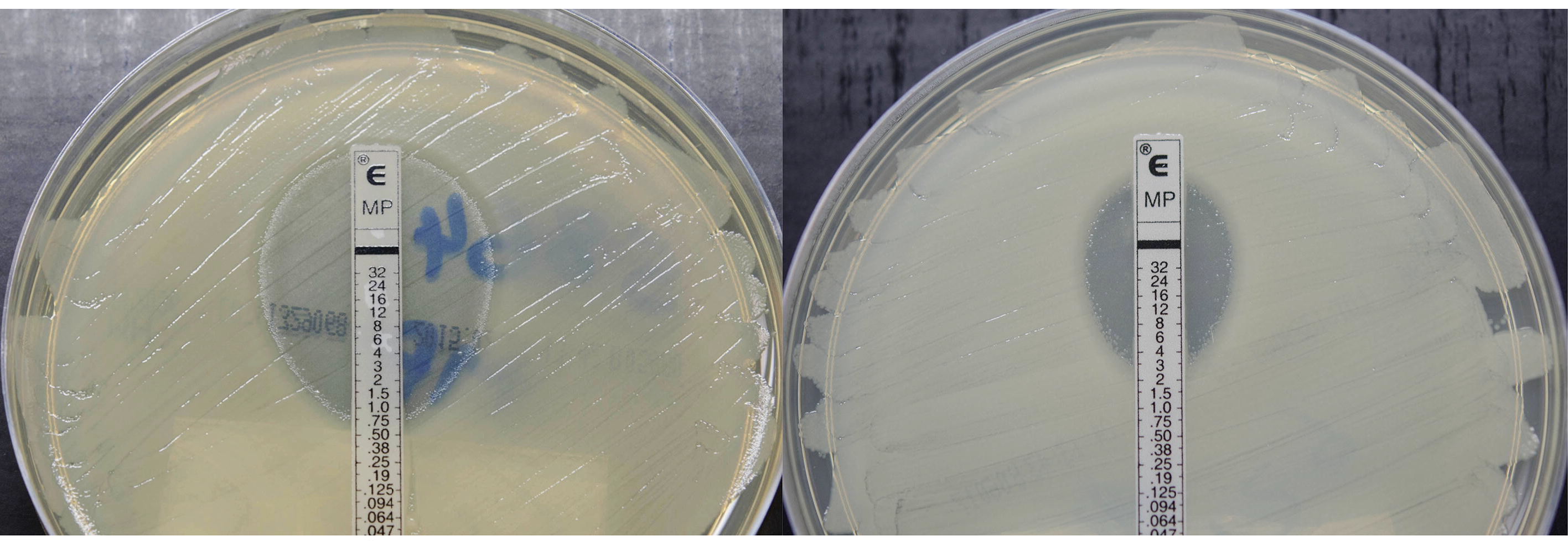
Representative meropenem Etest of a wild-type *B. pseudomallei* strain with a minimum inhibitory concentration (MIC) value of 0.75 μg/mL (left), compared to a strain that has developed reduced meropenem susceptibility with an MIC value of 3 μg/mL (right).

**Table 1:**
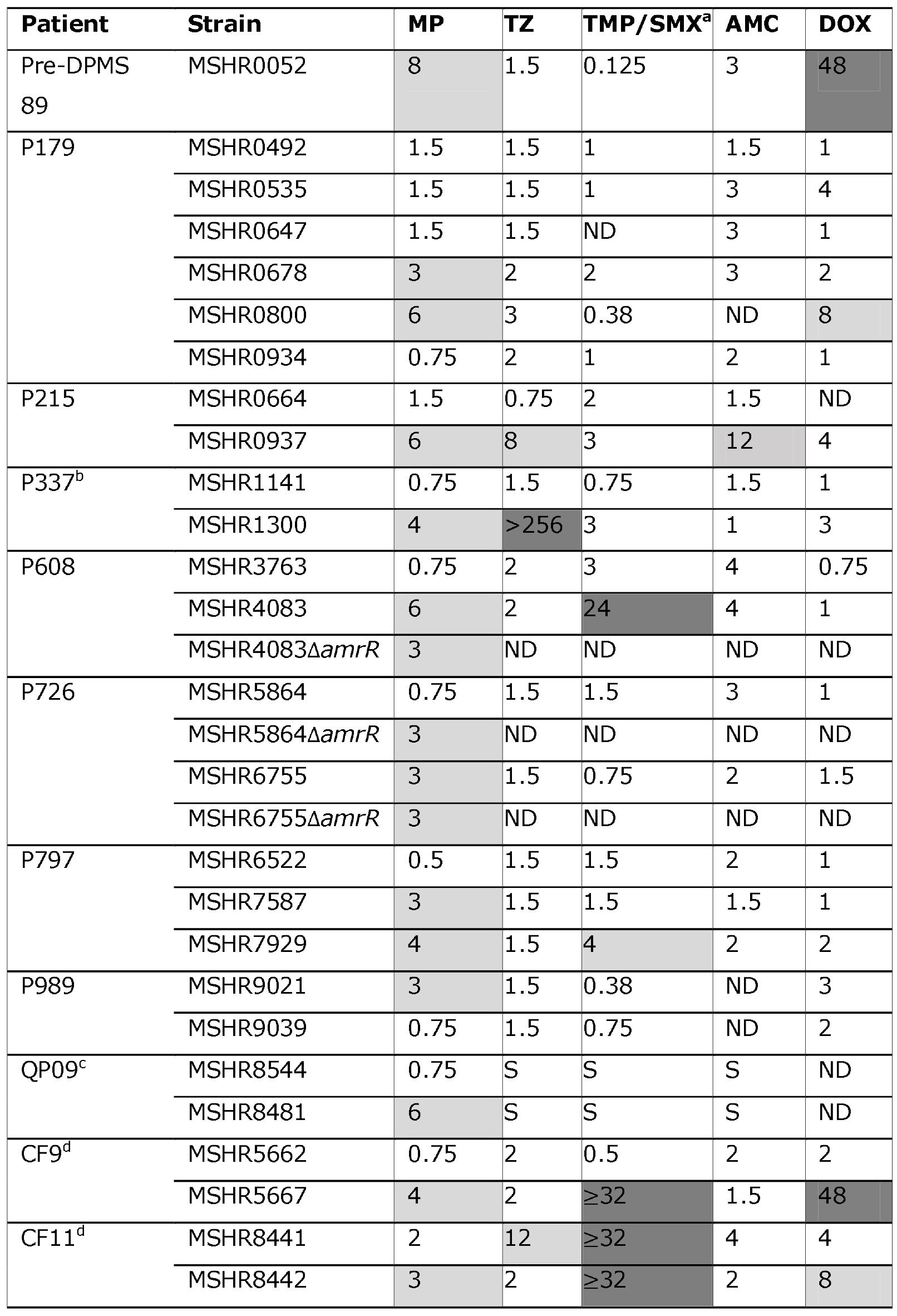
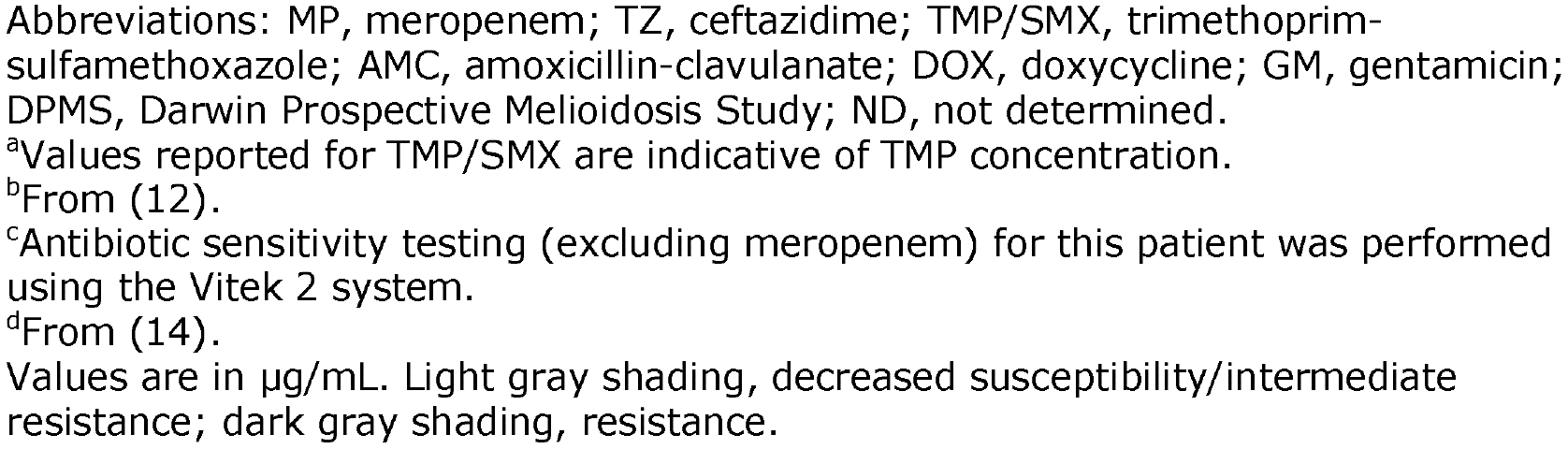
Minimum inhibitory concentration data for the *Burkholderia pseudomallei* isolates examined in this study.

| Patient | Strain | MP | TZ | TMP/SMX<sup>a</sup> | AMC | DOX |
| --- | --- | --- | --- | --- | --- | --- |
| Pre-DPMS<br>89 | MSHR0052 | 8 | 1.5 | 0.125 | 3 | 48 |
| P179 | MSHR0492 | 1.5 | 1.5 | 1 | 1.5 | 1 |
|  | MSHR0535 | 1.5 | 1.5 | 1 | 3 | 4 |
|  | MSHR0647 | 1.5 | 1.5 | ND | 3 | 1 |
|  | MSHR0678 | 3 | 2 | 2 | 3 | 2 |
|  | MSHR0800 | 6 | 3 | 0.38 | ND | 8 |
|  | MSHR0934 | 0.75 | 2 | 1 | 2 | 1 |
| P215 | MSHR0664 | 1.5 | 0.75 | 2 | 1.5 | ND |
|  | MSHR0937 | 6 | 8 | 3 | 12 | 4 |
| P337<sup>b</sup> | MSHR1141 | 0.75 | 1.5 | 0.75 | 1.5 | 1 |
|  | MSHR1300 | 4 | >256 | 3 | 1 | 3 |
| P608 | MSHR3763 | 0.75 | 2 | 3 | 4 | 0.75 |
|  | MSHR4083 | 6 | 2 | 24 | 4 | 1 |
|  | MSHR4083Δ*amrR* | 3 | ND | ND | ND | ND |
| P726 | MSHR5864 | 0.75 | 1.5 | 1.5 | 3 | 1 |
|  | MSHR5864Δ*amrR* | 3 | ND | ND | ND | ND |
|  | MSHR6755 | 3 | 1.5 | 0.75 | 2 | 1.5 |
|  | MSHR6755Δ*amrR* | 3 | ND | ND | ND | ND |
| P797 | MSHR6522 | 0.5 | 1.5 | 1.5 | 2 | 1 |
|  | MSHR7587 | 3 | 1.5 | 1.5 | 1.5 | 1 |
|  | MSHR7929 | 4 | 1.5 | 4 | 2 | 2 |
| P989 | MSHR9021 | 3 | 1.5 | 0.38 | ND | 3 |
|  | MSHR9039 | 0.75 | 1.5 | 0.75 | ND | 2 |
| QP09<sup>c</sup> | MSHR8544 | 0.75 | S | S | S | ND |
|  | MSHR8481 | 6 | S | S | S | ND |
| CF9<sup>d</sup> | MSHR5662 | 0.75 | 2 | 0.5 | 2 | 2 |
|  | MSHR5667 | 4 | 2 | ≥32 | 1.5 | 48 |
| CF11<sup>d</sup> | MSHR8441 | 2 | 12 | ≥32 | 4 | 4 |
|  | MSHR8442 | 3 | 2 | ≥32 | 2 | 8 |

Abbreviations: MP, meropenem; TZ, ceftazidime; TMP/SMX, trimethoprim-sulfamethoxazole; AMC, amoxicillin-clavulanate; DOX, doxycycline; GM, gentamicin; DPMS, Darwin Prospective Melioidosis Study; ND, not determined.
<sup>a</sup>Values reported for TMP/SMX are indicative of TMP concentration.
<sup>b</sup>From (12).
<sup>c</sup>Antibiotic sensitivity testing (excluding meropenem) for this patient was performed using the Vitek 2 system.
<sup>d</sup>From (14).
Values are in µg/mL. Light gray shading, decreased susceptibility/intermediate resistance; dark gray shading, resistance.

For all three cases, high-quality closed or close-to-closed assemblies of the isogenic strain pairs (21) enabled comprehensive identification of all genetic variants between the pairs (Table S1). A single gene, *amrR*, was mutated in all three latter isolates but not in the original isolates (Figure 2; Table 2; Table S1). AmrR is the local TetR-type regulator of the resistance-nodulation-division (RND) efflux pump AmrAB-OprA, which is responsible for intrinsic aminoglycoside and macrolide resistance in *B. pseudomallei* (22). The latter P608 isolate encodes an in-frame deletion of four amino acids (Alal53_Aspl56del). Similarly, the latter P726 isolate possesses an in-frame deletion of three amino acids (Val60_Cys63del), and two latter isolates from P797 encode a missense SNP that results in a Gly30Asp substitution. No structural alterations, large deletions or duplications were observed in the strain pairs, thereby ruling out other genetic variants in conferring decreased meropenem susceptibility.

**Figure 2:**
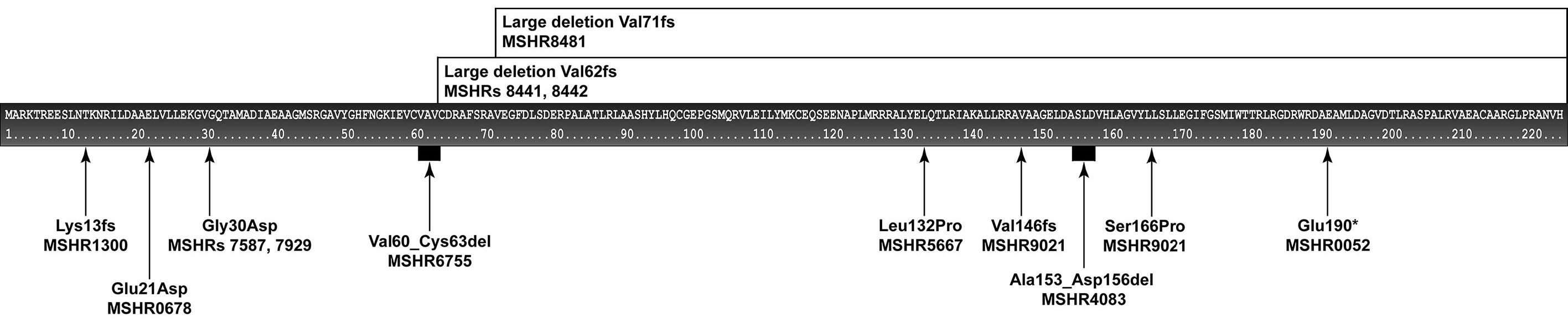
Distribution and annotation of mutations occurring within *amrR*. *Burkholderia pseudomallei* strains with decreased meropenem susceptibility usually harbor mutations in *amrR*, the regulator of the AmrAB-OprA resistance-nodulation-division efflux pump. Eleven distinct mutations were observed in *amrR*, with all mutations leading to a decrease in meropenem susceptibility. No overlap of mutations was observed for any of the strains, demonstrating that there are many possible mutations capable of inactivating or impairing AmrR function.

**Table 2:** Efflux pump regulator variants identified in Burkholderia pseudomallei isolates exhibiting decreased meropenem susceptibility.

| Patient | Strain | AmrR<br>(*BPSL1805*) | BpeR<br>(*BPSL0812*) | BpeT<br>(*BPSS0290*) |
| --- | --- | --- | --- | --- |
| Pre-DPMS P89 | MSHR0052 | Q190<sup>a</sup> | WT | WT |
| P179 | MSHR0678 | Q21D | WT | WT |
|  | MSHR0800 | WT | L85fs | WT |
| P215 | MSHR0937 | WT | D176A | WT |
| P337 | MSHR1300 | K13fs<sup>b</sup> | WT | S311R |
| P608 | MSHR4083 | A153_D156del<sup>c</sup> | WT | WT |
| P726 | MSHR6755 | V60_C63del<sup>c</sup> | WT | WT |
| P797 | MSHR7587 | G30D | WT | WT |
|  | MSHR7929 | G30D | WT | WT |
| P989* | MSHR9021 | S166P; A145fs<sup>d</sup> | WT | WT |
| Non DPMS QP09 | MSHR8481 | V71fs<sup>f</sup> | WT | WT |
| CF9 | MSHR5667 | L132P | WT | WT |
| CF11 | MSHR8441 | V62fs<sup>e</sup> | WT | WT |
|  | MSHR8442 | V62fs<sup>e</sup> | WT | WT |

Abbreviations: del, deletion; WT, wild-type; fs, frameshift
\*Isolates from this patient were intentionally sequenced from a potentially mixed population to capture population diversity. For MSHR9021, two AmrR mutants, S166P and A145fs, at ratios of ~66% and 33%, respectively (Figure S3). No AmrR mutants or mixtures were observed in the initial strain from this patient.
<sup>a</sup>Frameshift indel increases protein length from 223 to 288 residues
<sup>b</sup>Frameshift indel shortens protein length from 223 to 185 residues
<sup>c</sup>In-frame indel shortens protein length from 223 to 219 residues
<sup>d</sup>Frameshift indel shortens protein length from 223 to 183 residues
<sup>e</sup>Frameshift indel increases protein length from 223 to 285 residues
<sup>f</sup>Frameshift indel shortens protein length from 223 to 117 residues

### Functional characterization using scar-less knockouts and RT-qPCR verifies the role of *amrR* in upregulation of the RND efflux pump AmrAB-OprA

Scar-less *amrR* gene knockouts were created from two mutant isolates (MSHR4083; P608 and MSHR6755; P726) and one wild-type control strain (MSHR5864; P726). Removal of *amrR* from the three strains led to an MIC value of 3 μg/mL, regardless of the parental strain’s MIC. We then attempted *in trans* complementation with wild-type *amrR* using the pMo168 expression vector (23); however, due to these strains possessing very high kanamycin resistance following *amrR* removal, sufficient selective pressure needed to maintain pMo168 was not possible.

RT-qPCR was employed to determine whether *amrB* was upregulated in the strains exhibiting decreased meropenem susceptibility compared with their wild-type isogenic counterparts. Strains MSHR4083, MSHR3763, MSHR5864, MSHR6755, and their corresponding *amrR* knockouts (where available) were tested for increased *amrB* expression both with and without the presence of meropenem. For the wild-type strains, *amrB* expression was barely detectable regardless of meropenem presence or absence. All other strains with altered or deleted *amrR* showed significant up-regulation of *amrB* (range: 10 to 30-fold) in the presence of sub-inhibitory (0.25μg/mL) concentrations of meropenem, but no significant increased expression in the absence of meropenem.

### Additional strains with increased meropenem MICs also exhibit *amrR* mutations

Almost all primary isolates retrieved from the DPMS cohort have previously been screened for antibiotic resistance towards the clinically relevant drugs used in treatment (24). Latter strains from DPMS cases have generally only had repeat sensitivity testing if there has been a slow clinical response or recrudescent infection confirmed by culture (25). Through this testing, we identified a further four DPMS patients (P179, P215, P337, P989) harboring isolates that had developed elevated meropenem MICs over time. We also examined *B. pseudomallei* isolates obtained from four non-DPMS cases: two were cystic fibrosis (CF) patients with melioidosis with previously documented elevated meropenem MICs (14), case Pre-DPMS 89 was infected prior to the commencement of the DPMS, and case QP09 was from a different study site.

Comparative genomic analysis identified that, in all but two cases, isolates with decreased meropenem susceptibility encoded missense, frameshift or nonsense mutations in *amrR* (Figure 2, Table 2, Text S1 and Table S1). Curiously, two strains with decreased meropenem susceptibility were retrieved from P179, yet only one (MSHR0678) possessed an *amrR* mutation. The other strain, MSHR0800, instead encoded a frameshift mutation in the LysR-type regulator *bpeR* (*BPSL0812*; L85fs), the local transcriptional regulator of the BpeAB-OprB RND efflux pump (Table 2). A whole-genome SNP and indel phylogeny was constructed from all six isolates available from this patient and showed that these strains developed decreased meropenem susceptibility independently (Figure S1). A mutated *bpeR* was also observed in MSHR0937 from P215 (D176A), which otherwise did not have any mutations that could be linked to its decreased meropenem susceptibility (Table 2 and Text S1). In addition to having a truncated *amrR* (K13fs), MSHR1300 (P337) encoded a missense mutation in *bpeT* (*BPSS0290*; S311R), the local LysR-type regulator of the BpeEF-OprC RND-efflux pump. Interestingly, the latter isolate from P989 was a 66%/33% mixture of two strains that contained two independent mutations in *amrR*, a S166P substitution and A145 frameshift (Table 2, Figure S2 and Text S1).

### ReSistance towards other antibiotics in strains with decreased meropenem susceptibility

MIC determination of other clinically-relevant antibiotics identified multidrug resistance development in eight of the 11 cases, with acquired resistance or decreased susceptibility towards ceftazidime, TMP/SMX, amoxicillin-clavulanate or doxycycline having emerged in the latter strains (Table 1).

The most common co-resistance was towards TMP/SMX, which occurred in four cases. The latter isolates from P608, P797, CF9 and CF11 had elevated TMP/SMX MICs of 24, 4, ≥32 and ≥32 μg/mL, respectively, compared with parental MICs of 0.25-1 μg/mL. In the P608 mutant, a missense SNP affected *dut* (*BPSL0903*; also known as *dnaS*). This gene encodes for deoxyuridine 5′-triphosphate nucleotidohydrolase, an enzyme involved in pyrimidine metabolism. Although functionally determining the impact of this mutation was beyond the scope of this study, the Gly91Ala substitution likely contributes to high-level TMP/SMX resistance in this isolate as no additional mutations separated this isogenic pair. The P797 mutant encoded an in-frame deletion (Asp110_Gly116del) in a putative phosphoesterase (*BPSL2263*); this mutation was not present in the TMP/SMX-sensitive initial and midpoint isolates and was the only coding mutation that separated the latter and midpoint strains (Table S1). TMP/SMX resistance was also observed in both CF cases (14, 20). In CF11 (≥32 μg/mL), TMP/SMX resistance is most likely conferred by missense mutations affecting Ptr1 (R20-A22 insertion) and BpeS (the coregulator of BpeEF-OprC; V40I and R247L). The mechanism of TMP/SMX resistance in CF9 is not yet known, but may be attributed to missense SNPs in *dut* (N99S) or *metF* (N162P) (14) (Text S1).

Doxycycline resistance was observed in the *amrR* mutants from Pre-DPMS 89 and CF9 (48 μg/mL). We have recently shown that mutations affecting both AmrR and the SAM-dependent methyltransferase BPSL3085 lead to doxycycline resistance (26). Consistent with this prior work, the *amrR*-mutated strain from CF9 encodes a frameshift in *BPSL3085*, and in Pre-DPMS 89, two missense mutations affect this gene (Text S1). Strains from Pre-DPMS 89 and P215 also encode a PenA T147A variant, previously associated with high-level imipenem (32 μg/mL) and amoxicillin-clavulanate (32 μg/mL) resistance in the clinical *B. pseudomallei* strain, Bp1651 (18); however, despite encoding for this variant, all three strains (MSHR0052, MSHR0664 and MSHR0937) remained sensitive to imipenem (MICs = 0.5μg/mL), indicating that this mutation by itself does not confer imipenem resistance. It is possible that other mutations within *penA* are required to confer the high-level imipenem resistance phenotype observed previously (18). Interestingly, meropenem MICs showed no correlation with imipenem MICs, suggesting different pharmacodynamics of these two carbapenems in *B. pseudomallei*.

Decreased susceptibility towards ceftazidime was seen in P215 and CF11 (8-12 μg/mL), and high-level resistance (≥256 μg/mL) has previously been described in P337. A 10x duplication of a 36.7kb region encompassing *penA* (14), and a PenA C69Y mutant (12), are responsible for the elevated ceftazidime MICs in CF11 and P337, respectively. The molecular basis for decreased ceftazidime susceptibility in P215 is not yet known, although altered *penA* copy number and missense or promoter mutations can be ruled out, as can mutations in penicillin-binding protein 3 (*BPSS1219*). Based on genome comparison of the isogenic P215 pair, we propose that a deleterious frameshift mutation in *spoT*, the enzyme responsible for ppGpp metabolism (27), may sufficiently alter growth kinetics of the latter P215 strain, leading to CAZ resistance (Table S1).

## Discussion

Meropenem is a valuable antibiotic in the treatment of melioidosis, particularly for critically ill cases that require intensive care (28). The importance of meropenem is conferred in part by the fact that no primary resistance towards this antibiotic has ever been reported in *B. pseudomallei* (24). However, we have recently documented three cases whereby patients had extended and repeated episodes of blood culture positivity despite aggressive treatment with several rounds of antibiotics, including meropenem. A complicating feature of treatment was that the patients self-discharged from hospital multiple times, which greatly shortened their intended initial intravenous antibiotic regimens, based on the Royal Darwin Hospital melioidosis treatment duration protocol (29). All subsequently had difficulty clearing their *B. pseudomallei* infections and in two cases, they ultimately succumbed to their disease. Alarmingly, all were infected with *B. pseudomallei* strains that developed reduced sensitivity towards meropenem over the course of their treatment (MIC=3-8 *cf*. 0.5-0.75 μg/mL) (21).

Here, we expand upon this work by identifying and functionally characterizing the molecular basis for decreased meropenem susceptibility in *B. pseudomallei* isolates retrieved from 11 Australian melioidosis cases. In addition to the three previously reported cases, five new cases that were refractory to treatment were identified following a comprehensive review of ~1,000 patients enrolled in the DPMS, an ongoing study of melioidosis cases in the Northern Territory, Australia (5). Three additional cases not enrolled in the DPMS were also identified. A focus was placed on those patients with persistent culture positivity, or recrudescent or relapsing infections, and where longitudinal isolates were available for MIC testing and WGS. The eleven cases span 26 years and represent melioidosis patients living in two Australian states. The mutated strains were unrelated according to phylogenomic analysis (results not shown), demonstrating that the decreased meropenem susceptibility phenotype can potentially arise in any *B. pseudomallei* isolate given the right selective pressures.

Using a comparative genomics approach, we catalogued all genome-wide alterations (i.e. SNPs, small indels, duplications and larger deletions) separating the initial susceptible isolates from their mutated counterparts. Remarkably, isolates with decreased meropenem susceptibility from 10 of the 11 cases had accrued non-synonymous mutations within a single gene, *amrR*, which encodes AmrR, the TetR-type regulator of RND efflux pump AmrAB-OprA. In the remaining case, we observed an alteration in *bpeR*, the LysR-type regulator of the RND efflux pump BpeAB-OprB; this mutation was also observed in one of two isolates retrieved from P179 that encoded no mutations in *amrR* yet exhibited decreased meropenem susceptibility. One other patient, P337, harbored an isolate (MSHR1300) that, in addition to having a mutated *amrR*, encoded an altered *bpeT*, the LysR-type regulator of the RND efflux pump BpeEF-OprC. These results show that mutations affecting RND efflux pump regulation, and particularly AmrAB-OprA, cause increased MICs towards meropenem. Our findings provide a striking example of the value of comparative genomics in identifying the molecular basis for decreased antibiotic susceptibility and the contribution of convergent evolution in conferring a given phenotype.

To understand the link between RND efflux pump dysregulation and decreased meropenem susceptibility, we compared *amrB* expression levels in mutated and wild-type strains. The dramatic upregulation of *amrB* in the presence of meropenem and in the absence of a functional *amrR* (~10 to ~30-fold increase) is consistent with the regulatory role of AmrR on AmrAB-OprA, and our results show that meropenem probably acts as a substrate for efflux by this efflux pump. A similar phenomenon has also been documented in other Gram-negative pathogens. In *Pseudomonas aeruginosa* (30–32) and *Acinetobacter baumannii* (33, 34), over-expression of the MexAB-OprM and AdeABC efflux pumps, respectively, lead to decreased meropenem sensitivity although not high-level resistance. These studies also found that dysregulation of other efflux pump regulators can contribute to decreased meropenem susceptibility by increasing the expression of the other RND efflux pumps. Specifically, we observed three isolates with mutations in other efflux pump regulators (*bpeT* in MSHR1300 and *bpeR* in MSHR0800 and MSHR0937). Hayden and co-workers also identified a mutation within *bpeT* in *B. pseudomallei* 354e due to an 800kb inversion, increasing the meropenem MIC for this isolate to 6μg/mL (35). Taken together, these findings suggest that all three RND efflux pumps in *B. pseudomallei* can potentially efflux meropenem, although to what extent is not yet known. Alternately, they may regulate more than one RND efflux pump under certain conditions. The complex regulatory network controlling expression of these pumps is not yet well understood (36), and further work is needed to determine the precise roles of AmrR, BpeR and BpeT in the regulation of efflux pump expression in *B. pseudomallei*, including unraveling the mechanism underpinning the induction of AmrAB-OprA expression in the presence of meropenem. These questions are active areas of investigation in our laboratory.

Cross-resistance, a phenomenon whereby resistance towards one antibiotic can decrease susceptibility towards other antibiotics, has been described in many pathogenic bacterial species. One example is in methicillin-resistant *Staphylococcus aureus* (MRSA), an important cause of community-and hospital-acquired infections worldwide (37). The first-documented MRSA clones are thought to have become resistant to methicillin approximately 14 years before the first use of this antibiotic in clinical practice, suggesting that resistance was not a consequence of methicillin-driven selection but rather first-generation β-lactam use (38). In this vein, it is noteworthy that Pre-DPMS 89 and P337 harbored strains with reduced meropenem sensitivity yet were never treated with this drug, although they were both given doxycycline and P337 was also administered TMP/SMX (Text S1). These antibiotics are known substrates for at least one of the three *B. pseudomallei* RND efflux pumps (20, 26). In Pre-DPMS 89, the latter strain was doxycycline-resistant (MIC=48 μg/mL), and in P337, decreased TMP/SMX susceptibility (3 μg/mL) was seen (Table 1). Thus, we speculate that the increased doxycycline or TMP/SMX MICs led to cross-resistance towards meropenem in these cases. Interestingly, cross-resistance was not observed with imipenem, implying that these two carbapenems should be treated as separate drugs for the purposes of melioidosis treatment and isolate MIC testing; a substantial shift in current treatment dogma. The complex interplay of antibiotics and their cellular targets highlights the enigmatic role that multi-drug efflux systems can play in the development of antibiotic resistance in many bacterial pathogens, including *B. pseudomallei*. Further, our results show that resistance towards clinically relevant antibiotics can emerge during melioidosis treatment even when those antibiotics are not administered to the patient. A more intimate understanding of the mechanisms of cross-resistance in *B. pseudomallei* is needed to better identify and treat such mutants as they emerge under selection.

Current *B. pseudomallei* culture-based methodologies take between 24 and 72 hours for resistance determination, which can significantly delay the time in administering effective treatment (39). Predicting the resistance profile of a strain based on nucleic acid-based approaches is essential for the culture-independent detection of antibiotic resistance (40) and offers an attractive alternative to culture-based methods as a more rapid means for resistance detection. Due to the large number of mutations associated with decreased meropenem susceptibility, and the strong likelihood of novel mutations being uncovered that also contribute to this phenotype in other melioidosis cases, culture-independent platforms that can identify multiple mutations simultaneously will have the most impact and value in the clinical setting. Approaches such as portable, close-to-real-time next-generation sequencing of amplicons or whole bacterial genomes (e.g. Oxford Nanopore Technologies sequencing platforms (41)), or RT-qPCR assays that detect RND efflux pump upregulation, have the advantages of rapid turn-around-time, relatively low per-sample cost and broad target detection. We have preliminarily shown that *amrB* is dramatically upregulated (20-40-fold) in *amrR*-mutated *B. pseudomallei* strains that exhibit decreased meropenem susceptibility. Additional PCR assays targeting the BpeAB-OprB and BpeEF-OprC RND efflux pumps in *B. pseudomallei* may be useful in rapidly identifying *bpeR* and *bpeT* mutants, and we are currently evaluating a novel triplex RT-qPCR assay for this purpose (J. Webb *et al.*, manuscript in preparation).

Although the current study has identified and characterized mechanisms underpinning decreased meropenem susceptibility, there are still several unanswered questions that warrant further investigation. First, the precise clinical consequence of decreased meropenem susceptibility in *B. pseudomallei* is not yet clear. Our study focused on melioidosis cases that were persistently blood-culture positive, or that had recrudesced or relapsed, and excluded strains from acute, fatal infections or cases where eradication was achieved within the expected timeframe. It is possible that decreased meropenem susceptibility also occurs in these cases but remains undiagnosed. However, we deem this unlikely given that decreased meropenem susceptibility has only been identified on a handful of occasions despite extensive testing, suggesting that the decreased meropenem susceptibility phenotype is indeed correlated with poorer clinical response. Second, it is unclear whether the development of increased meropenem MICs would be grounds to exclude this drug as an available treatment option in all cases and indeed what alternative regimens may be appropriate. It is noteworthy that strains remained sensitive to imipenem despite efflux pump upregulation but co-resistance to ceftazidime, amoxicillin-clavulanate, doxycycline or TMP/SMX evolved in seven cases, suggesting that decreased susceptibility towards meropenem may be a risk factor for developing multidrug resistance due to synergism or cross-resistance.

## Conclusions

The carbapenem antibiotic meropenem is one of the most important drugs in the treatment of severe melioidosis cases. Here, we describe 11 cases where decreased meropenem susceptibility developed over the course of infection, with nine of the 11 patients having received meropenem as part of their treatment. Molecular characterization of all isolates exhibiting decreased meropenem susceptibility revealed several loss-of-function mutations leading to this phenotype, all of which up-regulated efflux pump expression, predominantly towards AmrAB-OprA, and to a lesser extent, BpeAB-OprB or BpeEF-OprC. Our study highlights the importance of treatment compliance and the need for early detection of emerging resistant populations. Continued assessment of treatment efficacy is essential for melioidosis management, especially in recrudescent or relapse cases, or when patients have difficulties clearing infection. Due to the large number of mutations leading to this phenotype, assays targeting efflux pump upregulation or rapid sequencing methods provide the most promising means of rapidly detecting emerging resistance.

## Materials and Methods

### Ethics statement

Ethics approval for this study has been approved as described elsewhere (5, 42).

### Clinical history and corresponding isolate description

Individual patient histories are detailed in Text S1. Unless otherwise stated, patients were enrolled in the DPMS. All underwent prolonged antibiotic treatment, often with multiple, repeated courses of antibiotics. Some patients took their own leave from hospital during treatment and several did so on more than one occasion.

### Culture conditions, DNA isolation and WGS

Unless stated otherwise, culture conditions, DNA extraction and WGS was performed as outlined previously (43).

### Meropenem MIC determination

Meropenem MICs were determined using Etests (bioMérieux) according to manufacturer’s instructions. The Clinical and Laboratory Standards Institute guidelines do not list MIC values for *B. pseudomallei* to meropenem, although there are guidelines for the related carbapenem antibiotic imipenem (≤4, 8 and ≥16 μg/mL for sensitive, intermediate and resistant, respectively). Based on our inhouse testing of >100 *B. pseudomallei* strains, we categorized decreased meropenem susceptibility as MICs ≥3 μg/mL. Guidelines for resistance cut-offs for other antibiotics were used as published in the CLSI guidelines and were as follows: ceftazidime and doxycycline S≤4, I=8 and R≥16 μg/mL, TMP/SMX S≤2/38 and R≥4/76 μg/mL, and amoxicillin-clavulanate S≤8/4, I=16/8 and R≥32/16 μg/mL.

### Reference genome assembly

The genomes of initial (meropenem-sensitive; MICs <2 μg/mL) and latter (decreased susceptibility; MICs ≥3 μg/mL) isolates from P608, P726 and P797 have previously been assembled to closure or near-closure using a combination of PacBio and Illumina data (21). These high-quality isogenic genomes were used to identify the molecular basis of decreased meropenem susceptibility. For other patients, only Illumina data were generated. Additional reference genomes were assembled to improved draft quality using MGAP v0.0.l (https://aithub.com/dsarov/MGAP---Microbial-Genome-Assembler-Pipelinel).

### Identification of molecular targets conferring decreased meropenem susceptibility

Comparative genomic analysis was performed using SPANDx v3.1.2 (44) to identify single-nucleotide polymorphisms (SNPs), small insertions/deletions (indels), large deletions and gene duplications between sensitive and decreased susceptibility isolates. SPANDx wraps Burrows Wheeler Aligner (45), SAMTools (45), the Genome Analysis Toolkit (GATK) (46), BEDTools (47) and SnpEff (48) into a single pipeline for ease-of-use and standardized outputs. The initial isolate from each patient was used as the reference genome for comparative genomic analysis; where this strain was unavailable, *B. pseudomallei* K96243 (49) was used for read mapping and variant annotation. Sequential isolates from the same patient were screened for rearrangements using Mauve v2.4.0 (50).

### Knockouts and complementation

*B. pseudomallei* scar-less gene knockouts were created using previously published methods (23). Primers used for vector construction and validation of *amrR* knockouts are described elsewhere (26). RT-qPCR of *amrB* expression was performed as previously detailed (26), with modifications. Primers and probe were redesigned and are as follows; amrB_forward 5′-TGTTCGCATGGGTGATCTCC-3′, amrB_reverse 5′-GACCGATTCCTCGACGACCT-3′ and amrB_probe 5′FAM-TGTTCATCATGCTGGGCGGCATC-3′ BHQ-1. Additionally, no pre-amplification of nucleic acid was required for detection and quantification. To measure the role of induction on *amrAB-oprA* expression, meropenem was included at the sub-inhibitory concentration of 0.25 μg/mL.

### Data availability

All WGS data is available under the following NCBI BioProject accession numbers: PRJNA272882, PRJNA300580, PRJNA393909, PRNJA397943 and PRJNA412120.

## Acknowledgements

We thank Vanessa Rigas and Barbara MacHunter (Menzies School of Health Research) for laboratory assistance, and the Pathology Department at Royal Darwin Hospital for sample collection. This work was funded by the National Health and Medical Research Council via awards 1046812 and 1098337. DSS is supported by an Advance Queensland Fellowship and EPP is supported by a University of the Sunshine Coast Fellowship.

